# Role of IL10 signaling in the circadian control of host response to Influenza infection

**DOI:** 10.1101/2025.03.03.641134

**Authors:** Kaitlyn Forrest, Martina Towers, Oindrila Paul, Hyeonbin Cho, Mahendra V. Padmini, Amruta Naik, Haeum Karrisa Lim, Yasmine Issah, Hitesh Deshmukh, Michael C. Abt, Kristin Hudock, Gregory R. Grant, Thomas G. Brooks, Shaon Sengupta

## Abstract

We have previously demonstrated that the circadian clock regulates the host response to influenza A virus (IAV) infection, conferring a time-of-day-specific protection –infection at dawn resulted in a threefold increase in survival and reduced immunopathology compared to infection at dusk. While IL10 is well-known for its immunoregulatory function, its role in IAV remains unclear, with studies reporting both protective and detrimental effects. Given the diurnal rhythmicity of IL-10 receptor (*Il10ra*) expression in the lung, we investigated the contribution of IL-10 signaling to time-of-day-specific IAV protection. We found that blocking IL10 signaling abrogated the time of day protection, leading to increased immunopathology characterized by enhanced lymphocyte infiltration and global immune activation (transcriptomic analysis). Interestingly, while later, IL-10R blockade also eliminated the time-of-day difference in IAV outcomes, it improved the outcome of dusk-infected mice. Furthermore, natural killer (NK) cell depletion suppressed IL-10 levels in bronchoalveolar lavage, suggesting a role for NK cells in regulating IL-10 signaling. In conclusion, incorporating the circadian context has not only clarified the IL-10 role in IAV infection but also underscored the pivotal influence of circadian regulation on immune responses.

## Introduction

Severe influenza infection is characterized by exaggerated and dysregulated inflammation^1,2^. Vaccines and anti-virals remain our first line in fighting influenza virus infection. However, our ability to target inflammation to limit pulmonary immunopathology remains limited. Circadian regulatory networks provide a mechanism through which the host can titrate the response to invading pathogens^3^, ensuring clearance while limiting immune-mediated pathology. IL10 is known for its immunoregulatory functions^4^ and is an attractive candidate for mediating protection from the influenza A virus (IAV). However, its role in IAV infection remains unclear. Higher IL10 levels were associated with worse outcomes in patients with severe respiratory infections. In mechanistic studies, one group has reported that loss of IL10 signaling by IL10R blockade worsened mortality from IAV, suggesting a protective role for IL10 in IAV^5^. However, another study showed that IL10^−/-^ mice had significantly lower mortality from IAV, implying that IL10 was detrimental to the host infected with IAV^6^. Notably, these studies did not consider the influence of the circadian clock on host responses. Building on our previous work^7^, we investigated whether IL10 signaling significantly contributes to the clock-controlled host protection from influenza A virus infection. We have previously shown that the circadian clock provides a time-of-day specific protection against IAV^7^. This was mediated not by higher viral replication early in the course of the infection but by a clock-driven immunoregulation that allowed the host to clear the virus with limited immunopathology. Our previous work has shown that differences in IL10 levels were associated with the time-of-day specific protection. While some have reported an association of circadian misalignment with altered levels of IL10^8,9^, the role of IL10 in the clock-driven host response to influenza infection has not been systematically examined before. This study tests the hypothesis that IL-10 is essential for circadian-mediated IAV protection.

## Results

### IL10 signaling is under clock control

We have previously shown that mice infected at ZT23 had threefold higher survival than mice infected at ZT11^7^ [ZT refers to Zeitgeber Time and is a way of specifying time with reference to the onset of light, the main zeitgeber; thus, ZT0 refers to the time when lights turn “ON” and ZT12 when lights go “OFF” in a 12 hr light-dark cycle]. This time-of-day specific protection was lost in mice wherein the clock was genetically disrupted by the loss of the core clock gene, *Bmal1*, thus confirming the role of the clock in mounting a host-protective response to IAV^7^. To test if early IL10-driven signaling contributes to the time-of-day specific protection from IAV, we decided to block IL10R. We define “early” as within the first 5 days post-infection (p.i.). IL-10R comprises the IL-10 receptor alpha (IL-10Ra) subunit, which binds the IL-10 ligand, and the IL-10 receptor beta (IL-10Rb) subunit; the latter is constitutively expressed on a large number of cells and shared across other cytokine signaling pathways. We chose to do this instead of IL10 neutralizing antibodies because we found that, as expected, *Il10ra* expression but not *Il10rb* showed an expression pattern that was suggestive of rhythmic oscillations on the publicly available dataset CircaDB^10^ [Fig 1A]. Identifying ZT6 and ZT18 as time points of maximal difference between gene expression, we tested the hypothesis that if *Il10ra* is under clock control, the diurnal difference between ZT6 and ZT18 should be abrogated in the lungs of *iBmal1*^*-/-*^ (*Bmal1*^*fl/fl*^*ERT2Cre*^+/-^) mice. *Il10ra* expression was highest at midday (ZT6) and lowest at midnight (ZT18). However, this difference was lost in *iBmal1*^*-/-*^ mice, suggesting IL10 signaling may be under circadian control [Fig 1B]. To test this definitively in the context of IAV infection, we infected age and sex-matched C57bl6/J mice at ZT23 or ZT11 with IAV (PR8; H1N1); we used well-validated anti-IL10R antibodies on days 0 (at the time of infection), 3, and 5 post-infection (p.i.) to block IL10 signaling [Fig 1C].

**Figure 1:**
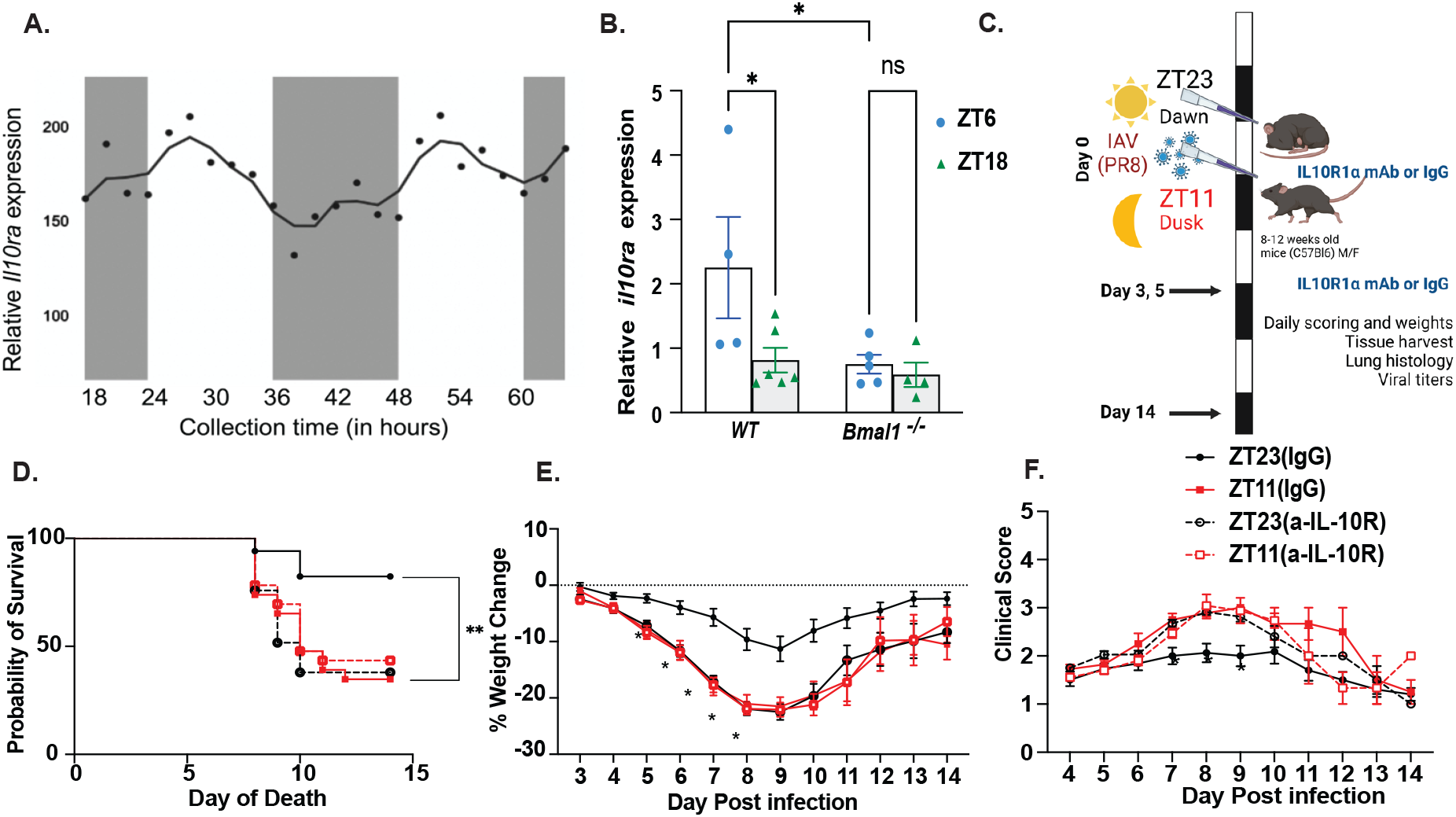
IL10 signaling is under clock control: (A) IL10 signaling is under clock control: (A) Expression of *Il10ra* appears rhythmic in naïve lungs (data extracted from publicly available dataset, CircaDB). q=0.233 by JTK. (B) Relative expression of Il10ra at ZT6 and ZT18 in *Bmal1*^*fl/fl*^ *ERT2Cre*^*+/-*^ and Cre^neg^ littermates. (C) Experimental design: two groups of mice maintained in 12 hr LD cycles were infected intranasally with IAV (40PFU) at either ZT23 (start of rest phase/dawn) or ZT11 (beginning of the active phase or dusk). A sub-group of these mice was treated with IL10R blocking antibody on days 0 (intranasal), 3, and 5 (intraperitoneal or i.p.). (D) Survival (n=17-29/group from 4 independent experiments). ****p<0.0001 by Log Rank test. (E) Weight loss (F) clinical scores (*p<0.05; n= 16-30/group by ANOVA for repeated measures with mixed-effect analyses with Dunnett’s correction for multiple comparison)

Interestingly, consistent with our previous work^7^, littermate controls receiving IgG maintained the time-of-day difference, with mice infected at ZT23 having significantly better survival than mice infected at ZT11 [Fig 1D; 82.3% survival in ZT23(IgG) controls versus 34.7% survival in ZT11(IgG)controls, p=0.00341 by Log-rank test and Suppl Fig 1]. However, in mice wherein IL10 signaling was blocked, the difference between ZT23 and ZT11 was abrogated and comparable to the ZT11 (IgG) group [37.9% survival in ZT23(a-IL-10R) versus 43.4% in ZT11(a-IL-10R)]. We posit that the conflict in the literature around the role of IL10 may be explained by considering the timing of the insult – no difference in the groups infected at ZT11, but a significant benefit from IL10 from the groups infected at ZT23 [Suppl Fig 1]. The weight loss trajectory also showed the time-of-day specific protection seen in the ZT23 IgG group was lost with the blockade of IL10 signaling (ZT23 (a-IL-10R)]) or ZT11(a-IL-10R) with outcomes comparable to that of the ZT11 (IgG) group [Fig 1E]. We also assessed the sickness behavior of the mice infected with influenza as a previously validated scoring system^7^ based on activity level, behavior, and respiratory distress (Supplementary Table 1). The ZT11 (IgG) had higher scores than the ZT23(IgG) group, consistent with higher morbidity in the former; IL10-blockade abrogated this difference with both antibody-treated groups showing more morbidity than the ZT23(IgG) group [Fig 1F].

### IL10 mediates its role in circadian protection from IAV by limiting immunopathology

To determine if IL10 blockade interfered with viral burden, thus leading to pathogen-mediated lung injury, we measured viral load in the lung tissue on days 5 and 8 p.i. Consistent with our previous findings, we did not see any significant difference in the viral load of IgG-treated mice. There were no differences in the groups with IL10R-antibody-treated mice infected at either ZT23 or ZT11 on day 5 post-infection (p.i.), which is also consistent with the literature^5^. By day 8, almost all mice from the four groups had cleared the virus [Fig 2A]. By day 8, on histology, we noted that the ZT11(IgG) mice sustained more injury than ZT23(IgG) as was evident on scoring based on (I) peri-bronchial inflammation, (ii) peri-vascular inflammation, (iii) alveolar inflammatory exudates, and (iv) epithelial sloughing/damage. Both ZT23(a-IL-10R) and ZT11(a-IL-10R) had similar lung injury, thus losing the time-of-day difference noted in the IgG-treated groups, and their severity was comparable to the ZT11 (IgG) group [Fig 2B]. However, even on day 5 p.i., when only 2 doses of the IL10-Rα antibody had been administered, and the viral titers were comparable across all four groups, hints of increased inflammation were beginning to surface in our analyses of the bronchoalveolar lavage (BAL) [Fig 2C]. The control mice infected at ZT23(IgG) had a lower total BAL cell count than the ZT11 group, consistent with our previous work. However, in the groups with IL10 signaling blockade, the BAL count in the ZT23 group was similar to that of the ZT11(a-IL-10R) group, overall comparable to the ZT11 (IgG) group [Fig 2C]. Overall, there were no differences in the distribution of the immune populations on day 5 p.i. [Suppl Fig 2A] Together, these results support the hypothesis that IL10 offers protection in the circadian gating of IAV-induced inflammatory response.

**Figure 2:**
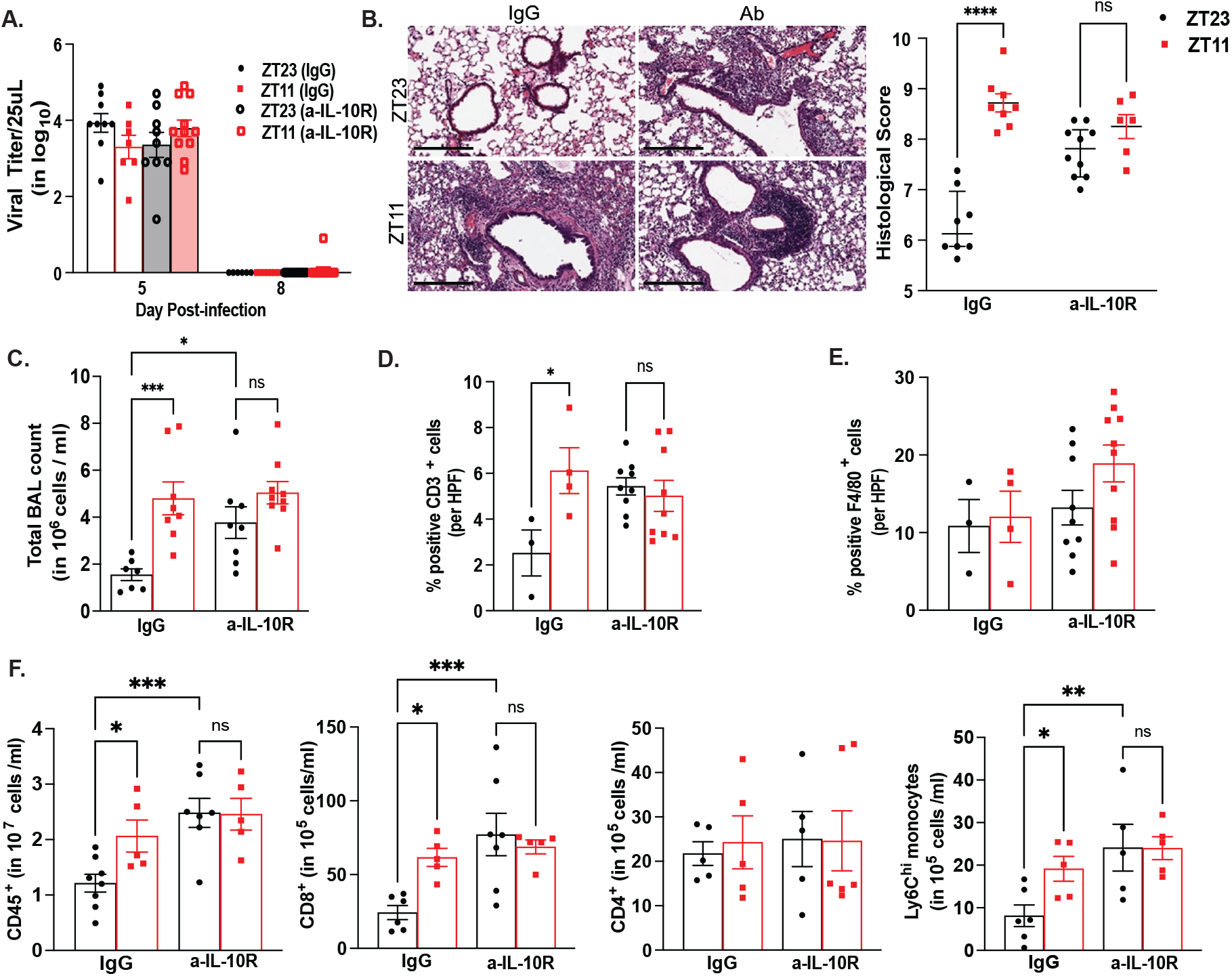
IL10 blockade affects immunopathology, but not viral clearance: Experimental design: After infecting mice with IAV at either ZT23 or ZT11 with or without IL10 signaling blockade, lungs were harvested for (A) Viral titers on days 5 and 8 p.i. (n= 6-9/group; 2-way ANOVA, p<0.0001 for day post-infection, N.S. for group and interaction, and (B) Histology. (right panel) Representative micrographs of the H&E-stained lung section from day 8 p.i. (scale bar =200μm). (Left panel): Severity of lung injury quantified using an objective histopathological scoring system by a researcher blinded to the study group (n=6-12/group; data as median and IQR; *p <0.05 by Wilcoxon rank-sum test) (C) Bronchoalveolar Lavage (BAL): Total cell count. (n=7-9/group; *p<0.05, ***p<0.0001 by 2-way ANOVA corrected for multiple testing). Representative lung sections with immunohistochemistry for (D) F4/80^+^ and (E) CD3^+^ cells in lung sections. (n=3-9/group; *p<0.05 by 2-way ANOVA corrected for multiple testing) (F) Flowcytometric elucidation of immune populations in mice infected at ZT23 or ZT11 with or without IL10 signaling blockade on day 8 p.i. (n=5-9/group). *p<0.05, **p<0.01, ***p<0.0001 by 2-way ANOVA corrected for multiple testing. Except for 2(B), data is expressed as mean ± SEM

### Early IL10 blockade is associated with the accumulation of lymphocytes

Next, we wanted to define what immune cell types contributed to the circadian protection mediated by IL10. Broadly, we enumerated myeloid (monocytes and macrophages) and T cells based on F4/80^+^ and CD3^+^ positivity, respectively, determining what cells contributed to peri-bronchial and peri-vascular infiltrates on lung histology in the groups wherein IL10 signaling was blocked [Fig 2B]. We noted more CD3^+^ cells in the ZT11 (IgG) lungs than the ZT23 (IgG) group. However, the total number of CD3^+^ cells was comparable in the ZT23(a-IL-10R) and ZT11(a-IL-10R) groups; overall, this was similar to the ZT11(IgG) group and significantly higher than the ZT23(IgG) group [Fig 2E]. In contrast, the total number of F4/80^+^ cells was comparable among the ZT23(IgG) and ZT11(IgG) groups. While the total number of F4/80^+^ cells in the IL10R-antibody treated groups was similar, interestingly, the ZT11(a-IL-10R) group had slightly more total F4/80^+^ cells on lung histology than the ZT23(IgG) groups [Fig 2E]. We further quantified the immune populations in the lung on flowcytometry and observed that there were more leukocytes in the ZT11(IgG), ZT23(a-IL-10R), and ZT11(a-IL-10R) groups than in the ZT23 (IgG) group [Fig 2F]. We found more CD8^+^ cells in the lungs of ZT11 (IgG) mice than ZT23 (IgG) mice. However, in both ZT23(a-IL-10R) and ZT11(a-IL-10R), there were more CD8^+^, but not CD4^+^ cells in the lungs on day 8 p.i., suggesting that the lymphoid population contributing to the immunopathology, as seen in Fig 2B, is driven by CD8^+^ cells [Fig 2F].

Interestingly, although the difference in the F4/80^+^ population of macrophages and monocytes was not as significant on lung histology [Fig 2E], there were more Ly6c^hi^ monocytes in ZT23(a-IL-10R) and ZT11(a-IL-10R) than in the IgG-treated groups [Fig 2F]. Since F4/80^+^ stains for a large group of myeloid cells, we concluded that IL10 blockade results in more inflammatory monocytes in the lungs that contribute to immunopathology. Overall, the loss of IL10 signaling results in more Ly6C^hi^ monocytes and CD8^+^ cells in the lungs.

### Blockade of IL10 signaling led to global immune activation in influenza infection

Next, we analyzed the transcriptional response to early IL-10 signaling after IAV infection to identify the pathways associated with optimal host outcomes and minimized immunopathology. To do so, we analyzed the transcriptome of whole lungs collected on day 8 p.i from the ZT23(IgG) and ZT23(a-IL-10R) groups (n=4-6/group including both sexes). We chose this time point to capture the effect of IL10 blockade adequately [the last dose of IL10R antibody was administered on day 5, Fig 1C] and provide a transcriptomic correlate for the phenotype described earlier [Fig 3A]. We found that 1711 genes were differentially expressed (DE) between the ZT23(a-IL-10R) and ZT23(IgG) groups with a >2 log fold change; of these, the expression of 524 genes was upregulated in the ZT23(IgG) group, and 1187 genes were upregulated in the ZT23(a-IL-10R) group respectively [Fig 3B]. Consistent with the increase in inflammatory monocytes in the ZT23(a-IL-10R) group, several genes including *Ccl4, Cxcl10, Ccl2, Isg15, Ly6c2, Oas3, Ccr5, Mx1, Il12rb1, Gzmb* and *Irf4* were upregulated in the ZT23(a-IL-10R) group compared to the ZT23(IgG) group [Fig 3C]. We have also noted many of these genes to be differentially regulated in our previous work comparing ZT11 and ZT23 lung transcriptome on day 6 p.i.^7^ Further, the two groups also differentially regulated many genes involved in cell cycle regulation (*Cdc20, Cdk1, Ccn4, Ccnd1*) and tissue remodeling (*Chil1, Chil3, Timp1*). Not unexpectedly, pathways implicated in IL10 signaling included those associated with innate immune activation and exaggerated inflammation, such as “pathogen-induced cytokine storm signaling pathways), B cell activation, Th1 pathway, and granulocyte and granulocyte diapedesis” as well as several pathways involved in cell proliferation. Further, based on the DE genes’ analyses, we identified a few key upstream regulators [Fig 3D] of the effect of IL10 on response to IAV. In comparing ZT23(a-IL-10R) versus ZT23(IgG), as expected, we saw that IL10RA and Stat6 were both inhibited, and several cytokines, including IL6 and IFNG, and transcriptional regulators like STAT3 and STAT1, were activated. Other activated upstream regulators include *Bhlhe40 (Dec1)*, a gene known for circadian and non-circadian functions. To validate the findings of increased immune activation, hypercytokinemia, and immune cell migration on transcriptomic analyses, we measured some key cytokines and chemokines in the BAL fluid harvested on day 8 p.i. We found that the levels of CCL2, IFNγ, Cxcl10, and IL6 were higher in the ZT23 (a-IL-10R) group than in the ZT23(IgG) group [Fig 3E]. Thus, early IL-10 blockade in IAV infection triggered excessive immune activation without lowering the viral burden, leading to a significant increase in mortality.

**Figure 3:**
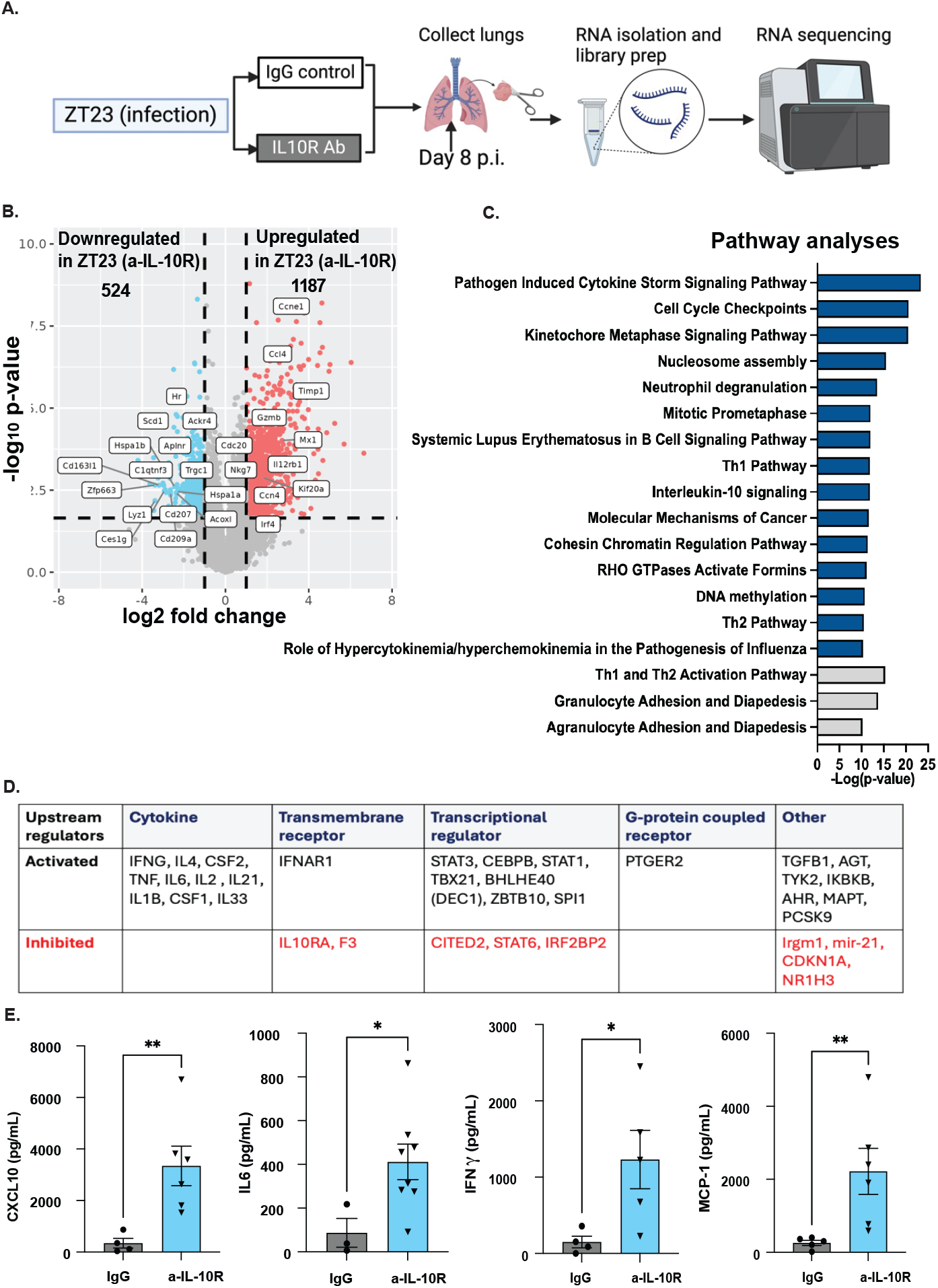
Blockade of IL10 leads to global immune activation following IAV infection. (A) Experimental design for RNA seq experiment comparing the transcriptome of mice infected at ZT23 with or without IL10blockade on day 8 p.i. (B) The plot of log-adjusted fold change for ZT23(IgG) and ZT23(Ab) showing the directionality of the most differentially expressed genes. (C)Top pathways based on the Ingenuity pathway analyses of DE genes between the two groups infected at ZT23. (D) Upstream regulators were identified based on the comparison of DE genes for ZT23(IgG) and ZT23(Ab) (E) Cytokine assay from homogenized lungs on day 8 p.i. *p<0.05, **p<0.01 by Mann-Whitney test.

### Transcriptional profiles underlying the effect of time-of-day and IL10 signaling

Blocking IL10 signaling phenocopies the time-of-day effect on histology, with the difference between the ZT23(IgG) and ZT11(IgG) groups being comparable to the ZT23(IgG) and ZT23 (a-IL-10R) group, as seen in Fig 2B. We next hypothesized that if IL10 signaling is one of the major pathways through which the circadian clock confers the time-of-day specific protection, the pathways and upstream regulators mediating the time-of-day difference between ZT23(IgG) and ZT11(IgG) should have considerable overlap with those mediating the effect of IL10 blockade seen in the comparison between the ZT23(IgG) and ZT23 (a-IL-10R) groups. Thus, we now compared the transcriptomic profile of three groups—ZT23(IgG), ZT23 (a-IL-10R), and ZT11(IgG) [Fig 4A]. As seen in Fig 4B, 847 genes were differentially expressed between the ZT23 (IgG) and ZT11 (IgG) groups with a >2 log fold change; of these, the expression of 259 genes was upregulated in the ZT23(IgG) group, and 588 genes were upregulated in the ZT11(IgG) group respectively. Of these 847 DE genes, more than half (492 genes) were shared with the DE genes from the ZT23 (IgG) versus ZT23(a-IL-10R) comparison above [Fig 3B]. Of these 492 common genes, 101 genes were upregulated in ZT23(IgG) group, and 389 genes were upregulated in both the ZT11(IgG) group and the ZT23(a-IL-10R) groups; only 2 genes (Spon2, Npas2) were differentially regulated in the opposite direction, being upregulated in ZT11(IgG) but downregulated in ZT23(a-IL-10R), both in comparison with the ZT23(IgG) group. Next, we drew three comparisons: ZT23 (IgG) versus ZT23(a-IL-10R); ZT23 (IgG) versus ZT11(IgG); and ZT23(a-IL-10R) versus ZT11(IgG). The 492 differentially expressed genes were well-represented across the two ZT23(IgG) comparisons [Fig 4B]. However, there were very few differentially expressed genes above the ≥2 log fold change in the ZT23(a-IL-10R) versus ZT11(IgG) comparison, suggesting that there was significant concordance in the mechanisms through which the circadian clock and IL10 protect and shape the host response to IAV. Interestingly, pathway analyses of these common DE genes included those associated with innate immune activation and amplification of immunopathology through adaptive immune response, such as “communication between adaptive and innate immune cells, B cell activation (as SLE signaling), pathogen-induced cytokine storm signaling pathways, IL15, Interferon signaling, and NFAT signaling.” Finally, as another check for mechanistic overlap between the IL10 and circadian pathways in acute response to IAV, we identified the top upstream regulators based on the common 492 genes [Fig 4D]. Interestingly, we saw that IL10RA was inhibited, while cytokines, IL6 and IFNγ, were activated along with STAT1 and TGFB1-these being conserved from our earlier comparison in Fig 3D. Overall, the overlap between these DE-expressed genes and the upstream regulators across these two comparisons supports the idea that IL10 signaling is an important contributor to the circadian control of host response to IAV.

**Figure 4:**
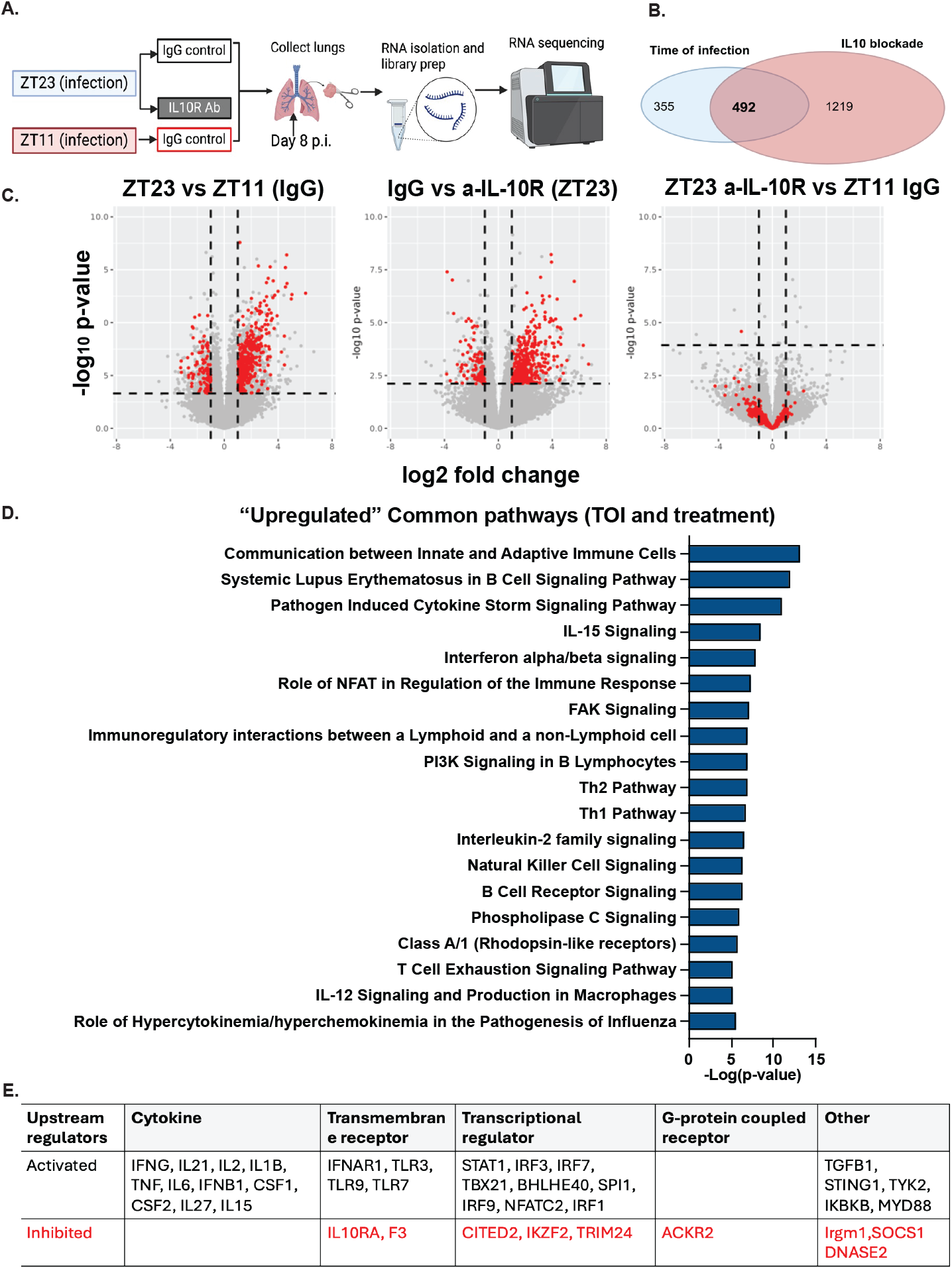
Significant overlap in the transcriptional profiles underlying the effect of time-of-day and IL10 signaling. (A) The experimental design for the RNA seq experiment comparing the transcriptome of mice infected at ZT23 with or without IL10blockade and ZT11 (IgG) on day 8 p.i. n= 4 for ZT23(IgG) and n=6 for ZT23(Ab), n=4 for ZT11 (IgG) (B) Venn diagram showing the overlap of the DE genes list for the two comparisons: ZT23 (IgG) versus ZT11(IgG) and ZT23(IgG) versus ZT23(Ab). (C) Volcano plots comparing the three groups (Red dots denote the 492 common DE genes) (D) Top pathways based on the 492 common genes across the comparison of the IL10 blockade and time-of-day effect by Ingenuity pathway analyses. (E) Upstream regulators were identified from the pathways analyses of the 492 shared genes.

### IL10 signaling contributes to optimal lung repair

We have previously shown that the effect of the circadian clock is apparent in lung repair following IAV infection^11^. To determine if the effect of IL10 signaling is apparent in recovering from lung injury following Influenza infection, we recovered the lungs to 14 p.i. While the ZT23(IgG) lungs had significantly lesser peri-vascular infiltrates, peri-bronchial infiltrates, better epithelial lining, and less epithelial hyperplasia, thus better lung architecture than the ZT11(IgG) group. This time-of-day specific protection was lost in the groups where IL10 signaling had been blocked [Fig 5A]. Next, we hypothesized that, given our transcriptomic profiling with the preponderance of a cell cycle control, tissue remodeling, and immunophenotyping results, the loss of IL10 signaling is likely associated with poor or dysplastic lung repair. In the absence of optimal proliferation of alveolar type 2 (AT2) cells to regenerate injured alveoli after IAV, aberrant expansion of basal cells results in dysplastic repair made by KRT5^+^ dysplastic pods. The ZT23(IgG) groups had minimal KRT5^+^ pods [Fig 5B; Supplemental Fig 3A], while the ZT11(IgG) had significantly more Krt5^+^ pods suggestive of dysplastic repair, consistent with our previous work. The time-of-day difference was lost in the ZT23(a-IL-10R) and ZT11(a-IL-10R) groups, both of which had more Krt5^+^ areas than the ZT23(IgG) group. Further, we noted that both on day 8 and day 14 p.i., the groups ZT23(a-IL-10R), ZT11(IgG), and ZT11(a-IL-10R) had significantly fewer Alevolar Type 2 (AT2) cells than the ZT23(IgG) group [Supplemental Fig 3B-C]. This suggests that a paucity of the AT2 cells in the dusk or IL10-blocked groups results in dysplastic regeneration. Thus, IL10 signaling is critical for the circadian clock-driven optimization of lung repair following IAV infection.

**Figure 5:**
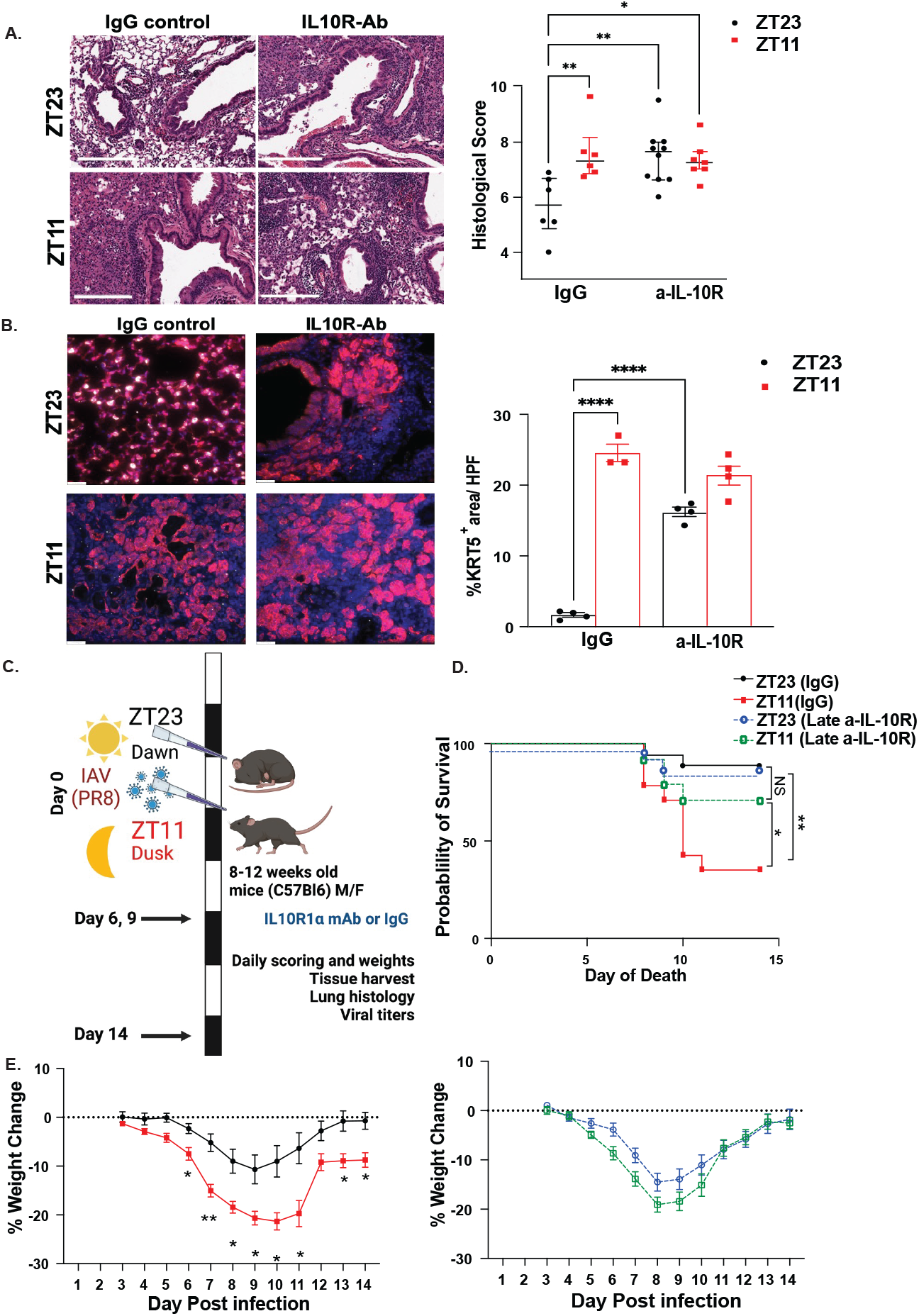
Late IL10 signaling blockade has a distinct effect on the circadian gating of lung injury after IAV infection: (A) Histology (Left panel) Representative micrograph of the H&E-stained lung section from day 14 p.i. (scale bar =200μm). (Right panel): Severity of lung injury quantified using an objective histopathological scoring system by a researcher blinded to the study group (n= 6-10/group; data as median and IQR; *p <0.05 by Mann-Whitney test). (B) Effect of IL10 blockade on lung repair characterized by representative image for KRT5 staining of lungs on day 14 p.i. (right panel) quantification of the KRT5+ areas. n=3-4/group from 2-3 independent experiments. (C) Experimental design for late IL10 -signaling blockade. two groups of mice maintained in 12 hr LD cycles were infected intranasally with IAV (40PFU) at either ZT23 or ZT11. A sub-group of these mice was treated with IL10R blocking antibody on days 6 and 9 (i.p.). (D) Survival *p<0.05, **p<0.001 by Log Rank test. (n=14-24/group from 4 independent experiments). ****p<0.0001 by Log Rank test (E) Weight loss. (n=13-20/group from 4 independent experiments) *p<0.05,**p<0.001 by mixed-effect analyses with correction for multiple testing.

### Late IL10 signaling blockade has a distinct effect on the circadian gating of lung injury after IAV infection

While our data reconciles the controversy in the field regarding the role of early IL10 in protecting against IAV by adding the circadian context, in several epidemiological studies in patients requiring invasive support due to respiratory failure associated with pulmonary viral infections, higher levels of IL10 in the lungs was associated with worse outcomes. Considered together, we hypothesized that the role of IL10 signaling in protecting against lung injury depends on the phase of injury-with early IL10 response being host-protective and late IL10 signaling deleterious to recovery. To test this, we infected mice at ZT11 or ZT23 as before; however, we blocked IL10 signaling using the IL10R antibody later after IAV infection [Fig 5C] on days 6 and 9 p.i. respectively. As expected, the ZT23(IgG) group continued to have a 3-fold higher survival than the ZT11 (IgG) group [Fig 5D; 89% survival in ZT23(IgG) versus 35% in ZT11(IgG); p<0.0001 by Log-rank test]. Unlike early IL10 signaling blockade, there was a minimal effect of blocking IL10 signaling late in the infection on the ZT23 infected groups, with the two groups having comparable survival [89% in ZT23(IgG) and 86% in ZT23(Late a-IL-10R) groups]. However, the late blockade of IL10 signaling conferred significant protection on the groups infected at ZT11 [71% survival in ZT11(Late a-IL-10R) groups versus 35% survival in the ZT11(IgG) group; p=0.0489 by Log-rank test]. Thus, as with early blockade, delayed IL-10 signaling blockade eliminated the time-of-day difference between the ZT23 and ZT11 groups; however, delayed blockade improved outcomes in the ZT11 group. Together, these data support our underlying premise that IL10 contributes to the clock-driven protection from IAV, but its role depends on the stage of influenza disease.

### NK cells contribute to IL10 signaling

In our transcriptomic analyses we noticed that NK signaling was differentially regulated pathways implicated in mediating the role of IL10 in clock-driven protection. Some other pathways, including IL12 and IL15 signaling, may also be involved in activating NK cells. Interestingly, we have previously reported that NK cell depletion early in infection abrogated the time-of-day difference in mortality from IAV^12^. Based on our data showing comparable results with IL10 signaling blockade, we hypothesized that NK cells contribute to the secretion of IL10 early after the infection. As in our previous work, we depleted NK cells one day before the infection and harvested BAL from infected mice 6 days p.i. [Fig 6A]. Consistent with our previous data, IL10 levels in the BAL of mice infected at ZT23 were significantly higher than those from the ZT11-infected mice. In the groups where NK cells had been depleted, IL10 levels were suppressed, irrespective of the time of infection, similar to the ZT11(IgG) group [Fig 6B]. In contrast to NK cells, interestingly, the loss of CD4 and CD8 cells on day 3 p.i. did not affect IL10 expression in the lungs of mice infected at ZT23 or ZT11 [Supplemental Figure 4]. Given the role of NK cells in mediating circadian protection from IAV through the production of IL10, we next hypothesized that the NK-depleted groups would phenocopy the effect seen with IL10 blockade. Indeed, we found that depletion of NK cells did not affect viral titers on days 2 and 6 p.i. [Fig 6C]. On day 6 p.i, lungs harvested from the ZT23(IgG) group had significantly less immunopathology than the ZT11(IgG), but in both NK-depleted groups infected at either ZT23 or ZT11, the injury was severe and similar to the ZT11(IgG) group [Figure 6D]. We chose day 6 for our analyses, given that this coincided with the difference in IL10 levels [Fig 6B]. These data support the hypothesis that NK cells mediate IL10 production; whether this is through direct secretion of IL10 by NK cells or indirectly by NK cells supporting the production of another subset of cells remains to be determined. Since clock proteins are transcriptional regulators, we next hypothesized that the clock regulates IL10 directly through transcriptional control of IL10 production. To test this, we undertook a chromatin immunoprecipitation (ChIP) assay using the naïve WT lungs. Using bioinformatic analyses, we found that the promoter region of IL10 contained at least one potential E-box motif (CACGTG) to which BMAL1 could be recruited. We then compared the fold enrichment of *Il10* and *Per2* (a known target of BMAL1 protein) following ChIP with anti-BMAL1 antibodies. Per2 showed approximately 6-fold enrichment of BMAL1 recruitment to its promoter over IgG control [Fig 6E]. IL10 was 20-fold enriched over its control, confirming that BMAL1 protein binds to the E-box motif in the IL10 promoter [Fig 6E]. Thus, we show that IL10 can serve as a target of the circadian clock via BMAL1. Considered together, the similarities in the effect of NK cell depletion and IL10 blockade support the idea that the NK cells are a critical regulatory innate immune population that mediates the clock gating of the host response to IAV through the IL10 signaling pathway and that the IL10 can be a direct transcriptional target in the context of acute inflammation.

**Figure 6:**
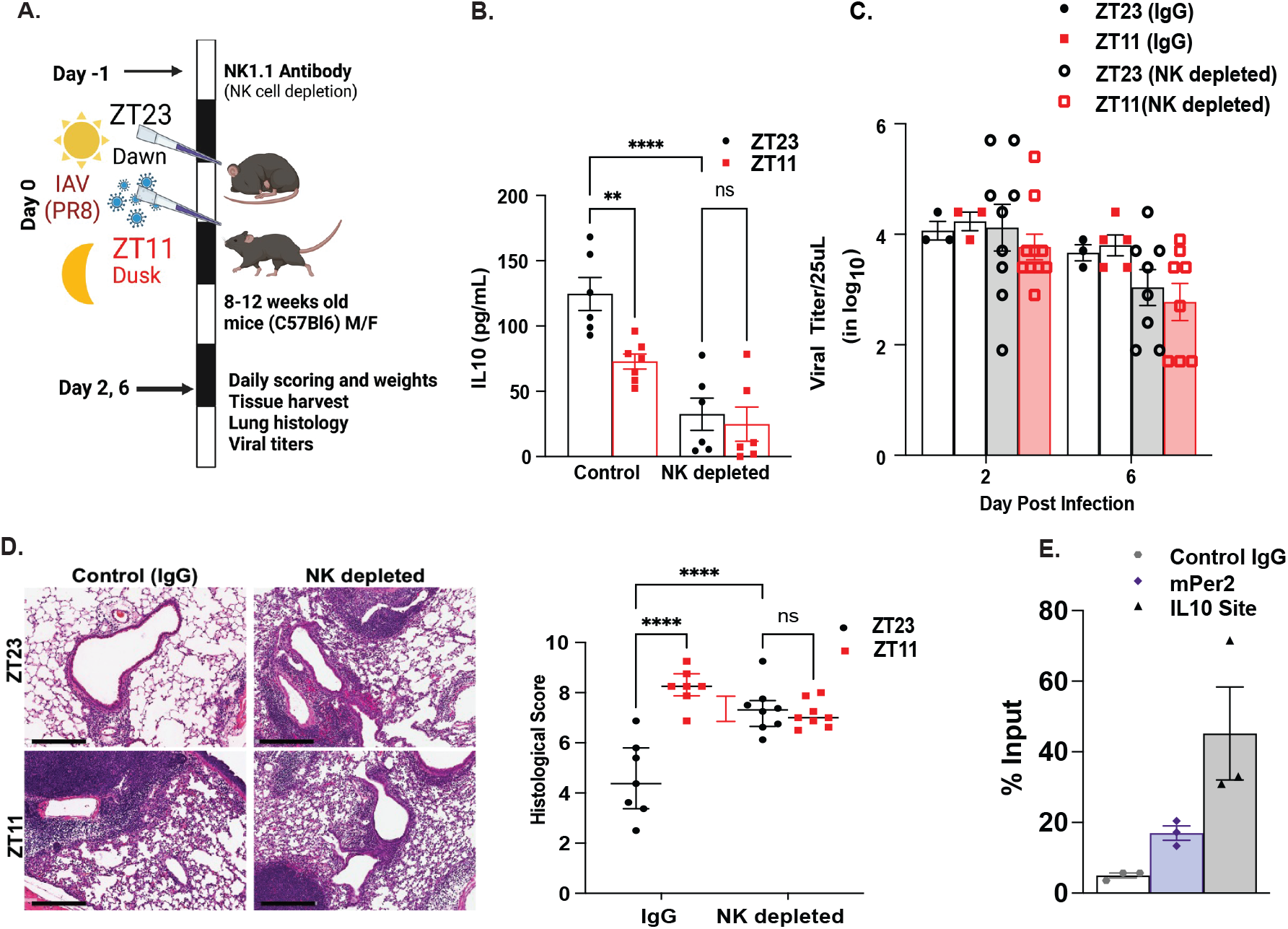
NK cells contribute to IL10-driven circadian protection from IAV: **(A)** Experimental scheme of NK cell depletion followed by IAV infection (B) IL10 levels in BAL from day 6 p.i. (C) Viral titers on days 2 and 6 p.i. (n= 6-9/group; 2-way ANOVA) (B) Histology. (right panel) Representative micrograph of the H&E stained lung section from day 6 p.i. (scale bar =200μm). (Left panel): Severity of lung injury quantified using an objective histopathological scoring system by a researcher blinded to the study group (n=6-12/group; data as median and IQR; *p <0.05 by 2 way ANOVA *p<0.05 for time-of-day effect, treatment and interaction; adjusted for multiple testing-****p<0.0001). (E) Chromatin immunoprecipitation (ChIP) assay for BMAL1 occupancy for the promoter region of *Il10*. mPer2 primers were used as positive controls for the analysis. Data expressed as a percentage of input level normalized to IgG control (n=3 from 2 independent experiments p<0.01 by Kruskal-Wallis test)

## Discussion

Here, we demonstrate that the IL10 signaling pathway is essential to clock-driven temporal protection from IAV throughout the disease trajectory. In early IAV, IL10 protects against mortality and immunopathology. Also, by considering the role of IL10 in circadian protection, we can offer a unifying paradigm for the conflicting reports on the role of IL10 in IAV. If we split the two groups based on the time of day at infection, we can see how IL10 may be beneficial for the dawn group but not so for the dusk group. [Supplemental Figure 1]. Further, we found that early IL10-signaling limits immune-mediated pathology while not affecting viral burden significantly. This is consistent with our previous work and with the role of the circadian clock in regulating immune response rather than being predominantly anti-viral in IAV infection^7^. IL10 blockade led to pulmonary infiltration by CD8^+^ cells and inflammatory monocytes, contributing to excessive immune-mediated pathology. Transcriptomic analyses of the two groups infected at dawn--ZT23(a-IL-10R) versus ZT23(IgG) groups showed that pathways that were consistent with the immunoregulatory role of IL10 that include negating the effects of IFNγ^13^, restricting the production of T_H_1-inducing cytokine IL12^14,15^, and limiting the secretion of pro-inflammatory cytokines by myeloid cells^16^. By several objective measures of injury, as determined by histology, lymphoid cell infiltration, accumulation of CD8^+^ cells, and transcriptomic analyses, there were significant similarities between the ZT23(a-IL-10R) and ZT11(IgG). Our transcriptomic analysis of ZT23(a-IL-10R) and ZT11(IgG) against ZT23(IgG) controls demonstrated that the observed similarities extended beyond broad phenotypic characteristics to shared mechanistic pathways. We observed contrasting effects of late IL-10 blockade on IAV outcomes: ZT23 outcomes were unaffected, but ZT11 outcomes improved, abrogating the time-of-day effect (as seen with early blockade; Fig. 1D), albeit with varying disease severity. This highlights the dynamic and context-dependent role of IL-10 in clock-driven IAV protection.

Considering the numerous immune and non-immune cell types capable of context-specific IL-10 production and the cytokine’s pleiotropic effects, we speculate that both IL-10 source and target cells are subject to circadian regulation. The observed oscillatory pattern of *Il10ra* expression in the lung and its significant reduction in circadian-disrupted (*Bmal1*^-/-^) mice supports the role of the circadian clock in modulating cellular responses to IL-10. However, following an acute infection, such as IAV, our data support the notion that the circadian control of the host response to IAV is more likely to be mediated by the production and action of IL10. Following acute injury, several circadian targets may stop oscillating, and others may start oscillating. For example, *Bmal1*^*-/-*^ peritoneal macrophages produced less IL10 but more IL6 than WT macrophages upon TLR4 stimulation^17^. Similar findings were reported in LysMcre^+^ mice infected with streptococcus pneumonia; however, in this instance, loss of the myeloid clock protected the host when infected at ZT12^18^. In another report, *Bhlhe40*, a gene with both circadian^19,20^ and non-circadian function^21^, was found to be a direct transcriptional regulator of IL10 in the Mycobacter*ium tuberculosis* model. Incidentally, *Bhlhe40* was one of the upstream regulators that were conserved across the comparison of the effect of IL10 signaling and time-of-day effects in our transcriptomic analyses here [Fig 3D and 4E]. While beyond the scope of our current work to investigate, given the role of *Bhlhe40* in switching immune cells from inflammatory to anti-inflammatory states^22^, we speculate that *Bhlhe40 (or Dec1)* may serve as a regulatory node for IL10 signaling.

As noted in our previous work^7^, the depletion of NK cells phenocopies the effect of early IL10 blockade, suggesting a role of the former in the circadian regulation of IL10 signaling. Further, NK cell-depleted groups, irrespective of the time of infection, had significantly lower IL10 levels in the BAL than the NK-sufficient group infected at ZT23 [Fig 6]. Our data strongly supports the role of the NK cells in controlling the clock-driven IL10 production. Thus, our data support two main possibilities linking NK cells with IL10: first, NK cells are the clock-driven source of IL10 early in infection, or second, NK cells support the production of IL10 by myeloid cells. Interestingly, Perona-Wright et al^23^. have shown that while NK cells can be an essential source of IL10 in systemic infection, they do not contribute significantly to IL10 in acute influenza infection. Since the time of day when mice were infected was not considered, this possibility cannot be excluded altogether. At the same time, loss of the myeloid clock was shown to suppress IL10 response in other studies^17,18^, raising the possibility that the role of the NK cells may be to support myeloid cells secreting IL10. Future work will be needed to determine whether NK cells produce IL10 production by NK cells or indirectly through NK-myeloid interactions.

The IL10-circadian connection is further reinforced by observations in shift workers in human studies known to have a circadian misalignment; circulating B cells isolated from rotating shift-working nurses produced less IL10 than non-shift working controls^8^. While shift workers represent the classic example of circadian misalignment leading to adverse health outcomes, circadian disruption is ubiquitous in modern society, thus underscoring the importance of studying circadian rhythms as an essential determinant of overall health. In our analyses of the UK biobank dataset (which contained <20% shift workers), we noted that robust circadian rhythms were associated with a lower risk for hospitalization^24^ and lower 30-day post-discharge mortality from severe respiratory infections^11^, underscoring the overall importance of circadian rhythms as an essential but often underappreciated driver of host response. We have demonstrated that IL10 shapes clock-driven acute inflammation [Fig 1-2] and later repair from IAV infection [Fig 5A-B]. This is consistent with our previous findings, where the clock was relevant for acute inflammation^7^ and chronic repair phases^11^. Beyond vaccines and anti-virals, both directed towards the pathogen, current therapeutic approaches fail to target the host’s immune response, which is often a significant determinant of outcomes in many respiratory infections. Thus, there is a critical need for immunotherapies^25^ that modulate the host response for optimal clearance and recovery. Adjusting the host’s circadian clock or downstream clock targets, such as IL10, is an attractive new avenue of investigation as we move into host-directed and personalized immunotherapy approaches. Finally, our data also suggest that both the time of day as well as the time after Influenza infection may be critical factors determining the efficacy of immunomodulatory therapies.

## Supporting information

Legends for Supplemental data

## Acknowledgments

We are grateful to Shivani Rawat and Paine Fleisher for their help with animal husbandry and general lab maintenance. This work was supported by NHLBI R01HL155934-01A1(SS) and NHLBI-R01HL147472 (SS). HC was supported by CURF. We are also grateful to Lawrence C Eisenlohr and Edward E Behrens for helpful discussions.

## Author Contributions

SS conceived the project; SS, KT, KH, MA and HD designed experiments; KF, OP, MT, HC, MVP, HC and SS performed experiments and collected data. SS, KF, OP, MT, HC, and TB analyzed data. GG and TB was involved in the analyses and interpretation of the transcriptomic data. KF, SS, and TB wrote the original draft, and KH, MA and GG helped with revisions. SS supervised all research activities.

## Data Availability

The RNA-Sc-seq data is available at the NCBI GEO accession number GSE287759. All other data is available as source data.

## Methods

### Mice

Adult C57b/6 mice 8-18 weeks in age were bred in-house or ordered from Jackson Laboratories. *Bmal1*^*fl/fl*^:*CAGG*^*creERt2/+*^ mice were bred in-house and treated with tamoxifen between 6-8 weeks of age to induce Bmal1 depletion. All mouse experiments were approved by the University of Pennsylvania and Children’s Hospital of Philadelphia Institutional Animal Care or the Children’s Hospital of Philadelphia (protocol ID 21-001405) and met the stipulations of the Guide for the Care and Use of Laboratory Animals.

### Infections

Mice were placed in circadian boxes 2-4 weeks before infection to allow for adjustment to the reverse 12-hour light/dark schedule. Under light isoflurane anesthesia, mice were intranasally infected with 30-35 PFU of mouse-adapted H1N1 (PR8) at either ZT11 or ZT23. To block IL-10 signaling, mice were treated with IL-10R monoclonal antibody (Clone 1B1.3A, BioXCell BE0050) or polyclonal IgG (Clone MOPC-21, BioXCell BE0083) during the infection. Mice were treated with 100ug/10uL of IL10R-MAb administered intranasally D0 in 10uL PBS, 150ug/100uL of IL10R Mab administered intraperitoneally on day 3 in 100uL of PBS, and 100ug/100uL of IL10R MAb administered i.p. D5. For late IL10 blockade, IL10R antibodies were administered i.p. on day 6 and 9 p.i. Controls received 10uL IgG intranasally or 100uL of PBS I.p. All antibiody administration was done at ZT23 or ZT11, based on the group.

In experiments with NK depletion, mice were treated with 200ug of NK1.1 monoclonal antibody (Clone PK136, BioXCell BE0036) or IgG control i.p. 24 hours before infection. In experiments with CD4 and CD8 cell depletion, mice were treated with 200ug each of CD4 monoclonal antibody (Clone GK1.5, BioXCell BE0003-1) and CD8a monoclonal antibody (Clone 2.43, BioXCell BE0061) in 100uL PBS or 200ug IgG i.p. on day 3 post-infection.

Weight and clinical observations were taken daily to monitor the progression of infection. Mice were clinically assessed on a scale of behavioral presentations ranging from 0-6. Mice were humanely euthanized when they reached 25% body weight loss and a behavioral score greater than or equal to 3 when they lost 30% or more of their body weight, or when a behavioral score greater than or equal to 4.

### BAL Cytology and Histology

The trachea was cannulated with 20G catheter (Surflo). Lungs were gently lavaged with 600L of PBS + protease inhibitor in four passes. Following centrifugation, the supernatant from the first pass was collected and used for further analysis. The cell pellets were combined and resuspended in 1mL PBS, counted using Nexcelcom cell counter, and cytospins were made. Staining was performed using Harleco Hemacolor Complete Stain Set (EMD Chemicals, 65044) according to the manufacturer’s instructions. Following CO2 asphyxiation, lungs were perfused through the right ventricle and instilled with 10% formalin at 20mm H_2_O through the trachea. Lungs were then paraffin-embedded and stained with H&E stain, CD3 and F4/80 as indicated. Slides were digitally scanned on Aperio CS-O and were viewed on Aperio ImageScope. For each sample, 8 representative sections were taken per slide. Images were scored based on the following criteria: (i) peri-bronchial infiltrate (ii) peri-vascular infiltrate (iii) alveolar infiltrate, and (iv) epithelial damage. Histological and cytospin scoring was done in a blinded fashion.

### Viral Titration

Viral burden was determined via hemagglutination of RBCs as previously described (Sengupta et al., 2019). Briefly, lungs were harvested and homogenized in PBS-gelatin (0.1%) and frozen for preservation. Influenza virus presence was measured by infecting MCDK cells (RRID:CVL_0422) with 1:10 dilutions of lung homogenates at 37°C. After 1 hour of infections, 175uL of media (2ug/mL trypsin) was added and the cells were further incubated for 72 hours at 37°C. 50uL of supernatant was used to test the hemagglutination of turkey RBCs, which would indicate the presence of virus particles.

### qPCR Analysis

RNA extraction from the lungs was performed using TRIzol (Life Technologies). The RNA was purified using the RNeasy Mini Elute Cleanup Kit (Qiagen) according to the manufacturer’s instructions. The quantity and quality of RNA were determined on a Nanodrop One C UV-Vis Spectrophotometer (Thermofisher Scientific) and cDNA prepared with TaqMan. SYBR Green and TaqMan gene expression assays were used to measure mRNA levels of genes of interest, with eukaryotic 28S (Sigma) and 18S rRNA (Life Technologies), respectively, used as internal controls. Assays were run on QuantStudio5 Real-Time PCR system (Applied Biosciences), and the relative ratio of the expression of each gene was calculated using the 2^ΔΔCt^ method.

### Flow Cytometry

Following harvest, lungs were digested with DNAse and Liberase in Dissociation Media (DMEM + 2% FBS + 1% Pen/Strep + 1% L-Glutamine + 0.1% 2-mercaptoethanol) in 37°C shaking water bath for 20 minutes. Spleens were placed in 1mL of Dissociation Media Dissociated tissue was passed through a 70um cell strainer and treated with RBC lysis buffer, washed, and then resuspended in 1mL PBS + 2% FBS + 0.1% Sodium azide. Then 3×10^6^ cells were treated with Fc block for 5 minutes, washed and stained with indicated antibodies on ice and protected from light for 20-30 minutes. Flow cytometric data was acquired on FACS Cantos flow cytometer and analyzed on FlowJo software. All cells were pre-gated as live cells.

### Chromatin Immunoprecipitation (ChIP)

Lung samples were minced into small pieces and cross-linked with 1% formaldehyde to incubate for 10 minutes at room temperature on tube revolver. To stop cross-linking, 125mM glycine (Thermofisher Scientific) was added and samples were further incubated for 5 minutes before homogenized for 10s at 0.6m/s on MP FastPrep24 (MP Biomedical). Following washing with cold PBS and centrifugation, cell pellet was resuspended in swelling buffer (components) on ice for 30 minutes. Then, the crude nuclear preparation was centrifuged, and the nuclei resuspended in nuclear lysis buffer before sonicated for 35 minutes at 4°C on Diagenode BioRupter to an average size 300-500 base pair. Sonicated samples were diluted 10-fold in IP buffer (components) and then incubated with 5ug Bmal1 antibody (Abcam AB3350) or rabbit IgG control (Millipore CS200581) overnight at 4°C on tube revolver. DNA complexes were collected on Dynabeads A (Thermofisher Scientific, 1001D) and washed with dialysis buffer (components), three times with IP wash buffer(components), and then once with TE buffer (components) before eluted with elution buffer (components). Crosslinking was reversed via incubation with 0.3M NaCl at 65°C overnight, followed by proteinase K treatment (components) and DNA cleanup. qPCR was performed using hPer2 and IL10 primers on QuantStuido5. Data was analyzed using % input method.

### RNA-seq analyses

We quantified RNA-seq reads using Salmon v1.9.0 with the GRCm38 Ensembl v102 reference transcriptome^26^. We assessed ribosomal content using PORT v0.8.5e (https://github.com/itmat/Normalization) and discarded two samples from further analysis due to extreme ribosomal content (>80%). The two comparisons of interest were, first, ZT23(IgG) versus ZT23 (IL10R a-IL-10R), and second, ZT11(IgG) versus ZT23 (IgG). Gene-level quantifications were imported using tximeta v1.20.3^27^ and differential expression was run with DESeq2 v1.42.0^28^. A sex term was included in the DESeq model. To compare ZT11(IgG) versus ZT23 (IgG), we further controlled that zeitgeber time of collection was equal to that time of infection and, therefore, differed between the two conditions of interest. We used a separate RNA-seq experiment that included tissues collected at ZT0, ZT12, and ZT24, all without infection or treatment. The same pipeline also quantified this dataset and the two datasets were concatenated and passed to DESeq. This model then included terms for time of infection (a categorical value of either ZT11, ZT23, or NA for the dataset that included no infection), time of collection (where ZT0, ZT23, and ZT24 were all considered the same, and ZT11 and ZT12 were supposed to be the same), and sex. No interaction terms were included. The batch effect arising from the two datasets was controlled for only because the time of infection variable completely separated the two variables. The result of interest was then ZT11 versus ZT23 time of infection. Including the time of collection factor allowed DESeq to account for the expected circadian transcriptional variation from time of day.

