## Supplementary material for "Role of IL10 signaling in the circadian control of host response to Influenza infection": Legends for Supplemental data

Legends for supplemental figures

Table 1: Clinical severity scoring for morbidity

Figure 1: Survival data from early blockade of IL10-signaling as in Fig 1, stratified by the time of infection.

Figure 2: (A) Differential of the total Bronchoalveolar lavage (BAL) count. (B) Representative lung sections of the immunohistochemistry for CD3 and F4/80 on day 8 p.i.

Figure 3: (A) Representative images of lungs harvested on day 14 p.i. Lung repair was quantified by staining for KRT5(dysplastic areas) and LAMP3(Alveolar Type 2, AT2 cells). AT2 cells, quantified as LAMP3^+^ cells/HPF (n=3-4/group from 2-3 independent experiments). (C) AT2 cells quantified as above on day 8 p.i. (n=3-4/group from 2-3 independent experiments). *p<0.05, **p<0.01, ****p<0.00001 by 2-way ANOVA, corrected for multiple testing.
