## Supplemental data for "Role of IL10 signaling in the circadian control of host response to Influenza infection"

### Slide 1
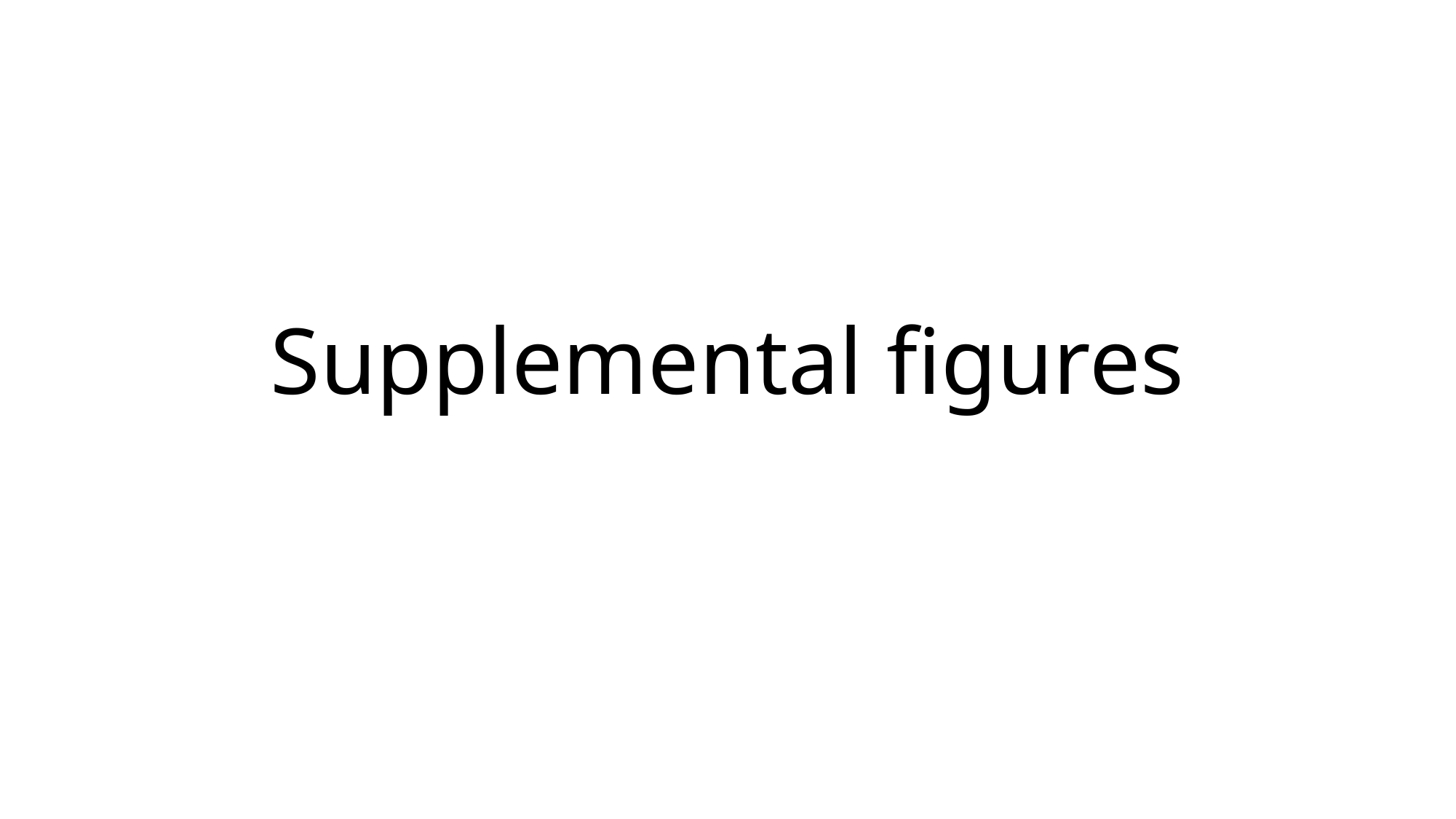

#### Supplemental figures

### Slide 2
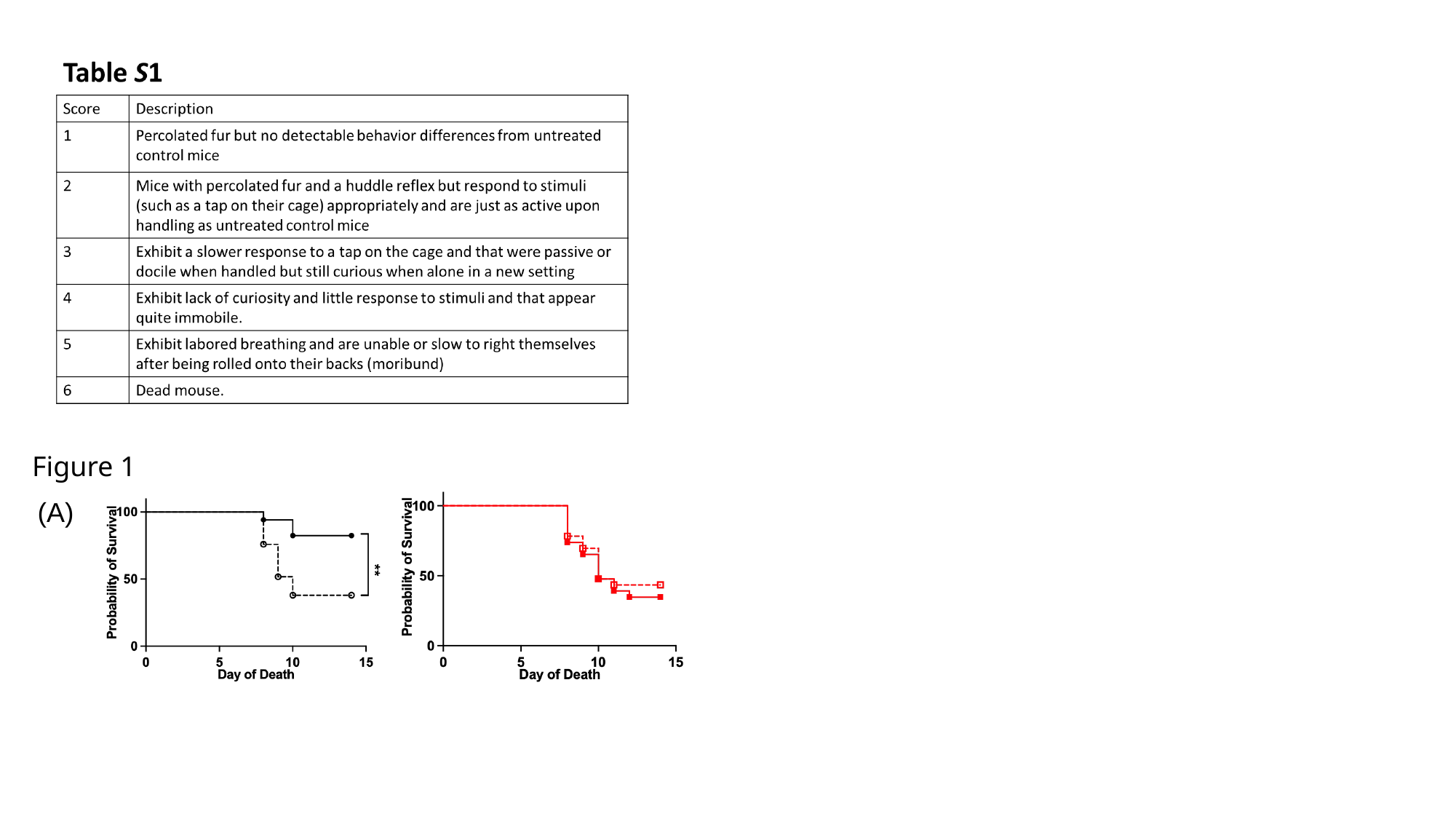

Figure 1
(A)

### Slide 3
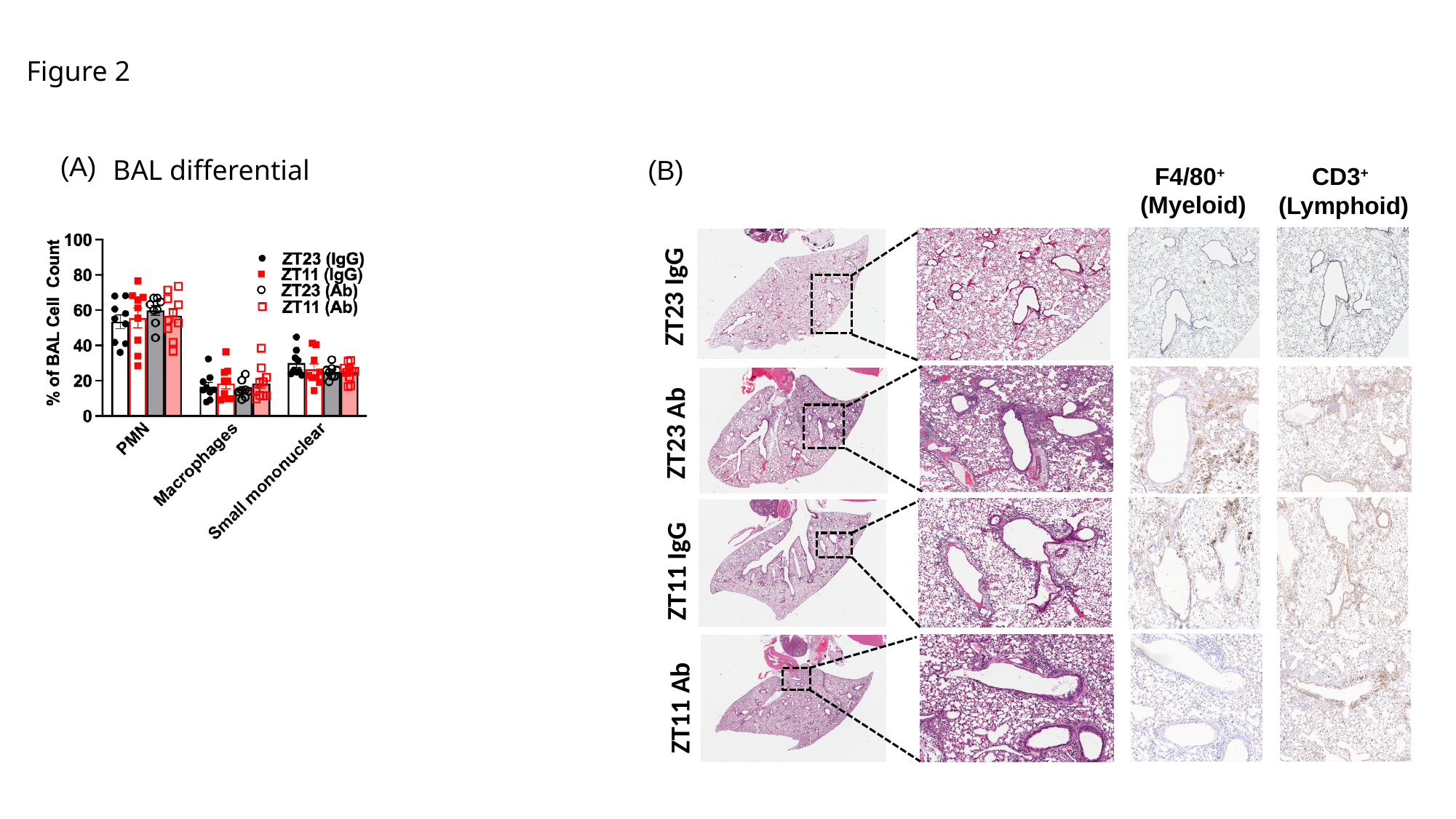

Figure 2
(A)
BAL differential
(B)
F4/80^+^
(Myeloid)
CD3^+^
(Lymphoid)
ZT23 IgG
ZT23 Ab
ZT11 IgG
ZT11 Ab

### Slide 4
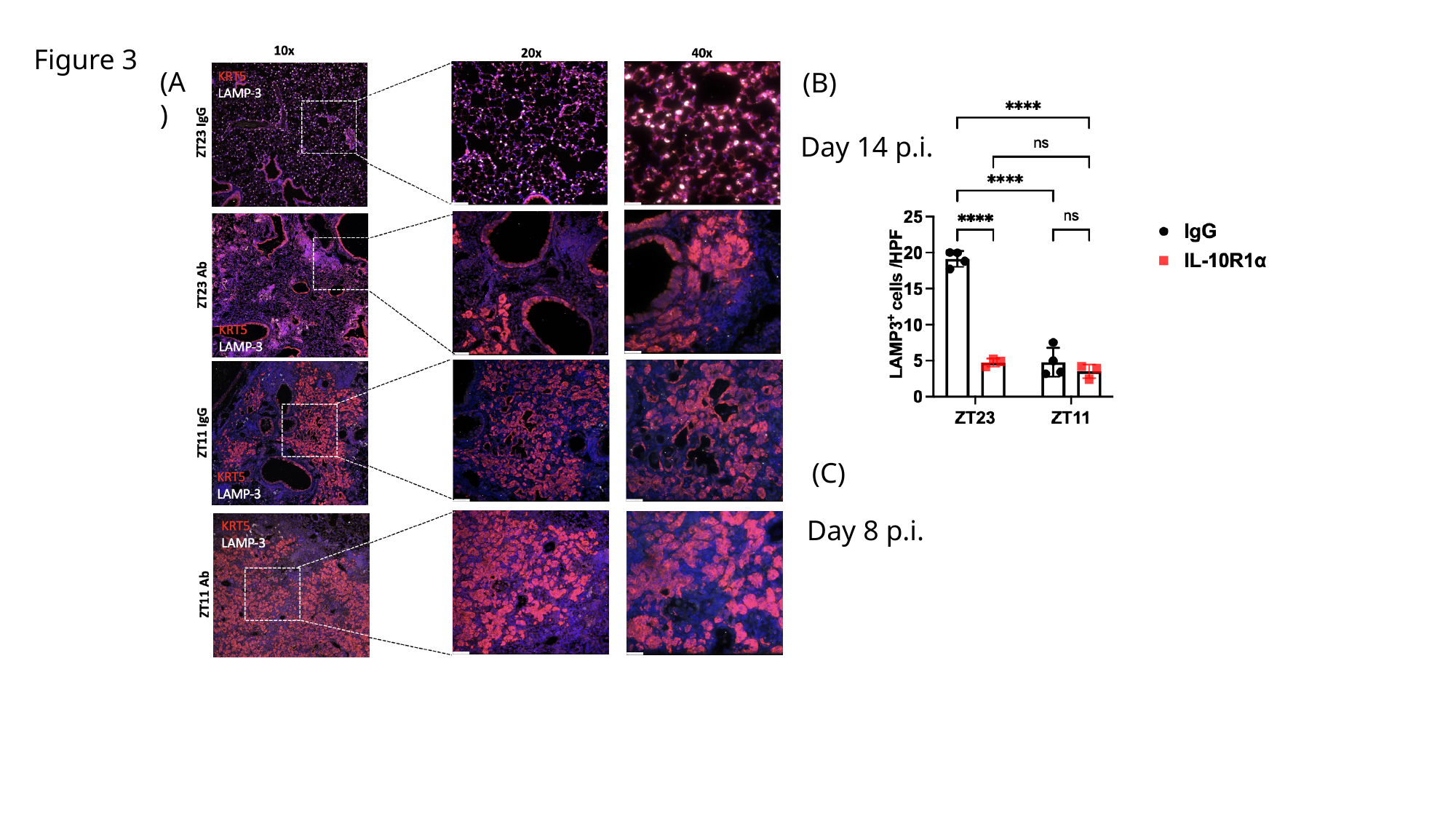

Figure 3
(A)
(B)
Day 14 p.i.
(C)
Day 8 p.i.

### Slide 5
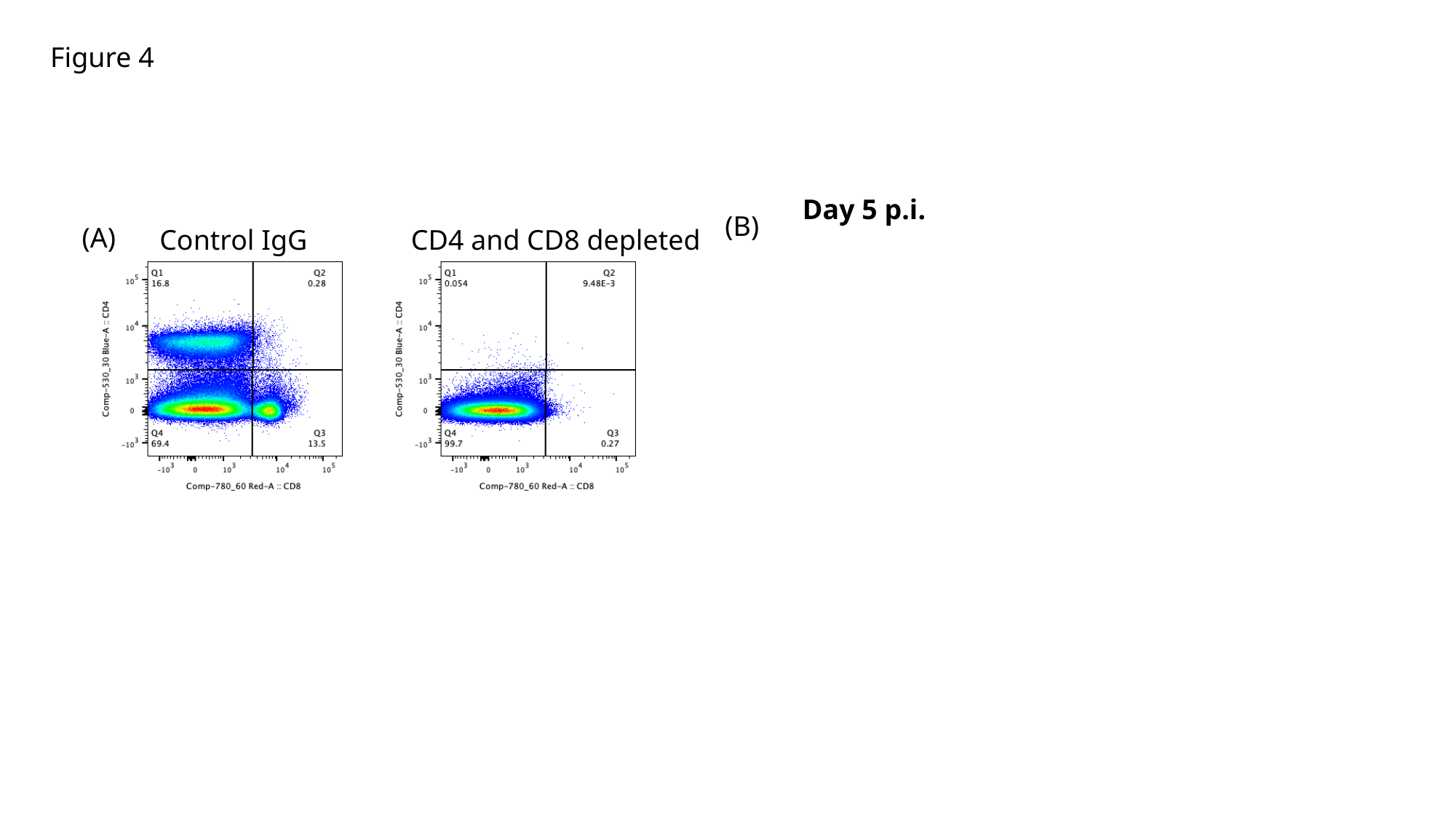

Figure 4
Day 5 p.i.
(B)
(A)
Control IgG
CD4 and CD8 depleted
